# Awake reactivation and suppression after brief task exposure depend on task novelty

**DOI:** 10.1101/809574

**Authors:** Ji Won Bang, Dobromir Rahnev

## Abstract

Previously learned information is known to be reactivated during periods of quiet wakefulness and such awake reactivation is considered to be a key mechanism for memory consolidation. We recently demonstrated that feature-specific awake reactivation occurs in early visual cortex immediately after extensive visual training on a novel task. To understand the exact role of awake reactivation, here we investigated whether such reactivation depends specifically on the task novelty. Subjects completed a brief visual task that was either novel or extensively trained on previous days. Replicating our previous results, we found that awake reactivation occurs for the novel task even after a brief learning period. Surprisingly, however, brief exposure to the extensively trained task led to “awake suppression” such that neural activity immediately after the exposure diverged from the pattern for the trained task. Further, subjects who had greater performance improvement showed stronger awake suppression. These results suggest that the brain operates different post-task processing depending on prior visual training.

## Introduction

The human brain has a remarkable ability to learn. A large number of studies have shown that post-learning neural processes play a critical role in consolidating new memories (Diekelmann & Born, 2010; McGaugh, 2000; Sasaki, Nanez, & Watanabe, 2010). One of the consolidation processes is the reemergence of the brain activities involved in the initial task during following wakefulness and sleep (Foster, 2017; Pfeiffer, 2017; Tambini & Davachi, 2019).

Awake reactivation, which refers to the reinstatement of previous learning during following rest was demonstrated in human neuroimaging studies. Previous studies found awake reactivation in the medial temporal lobe (Staresina, Alink, Kriegeskorte, & Henson, 2013; Tambini & Davachi, 2013), higher-order areas (Chelaru et al., 2016; de Voogd, Fernandez, & Hermans, 2016; Deuker et al., 2013; Guidotti, Del Gratta, Baldassarre, Romani, & Corbetta, 2015; Schlichting & Preston, 2014), and even the primary visual cortex (Bang, Sasaki, Watanabe, & Rahnev, 2018) after learning on various visual stimuli. The stimuli ranged from complex items, such as cartoon movies and animals to simple Gabor patches. This line of studies suggests that reactivation is likely to occur across most brain areas with most learning paradigms.

Although much work has focused on identifying awake reactivation after various types of learning, relatively little is known about which factors affect awake reactivation. Identifying influential factors is critical because it sheds a light on how the brain selects which information becomes reactivated. Motivated by this question, we focused on the effect of the novelty of the task on awake reactivation.

In order to investigate how the novelty of the task affects awake reactivation, we trained subjects on one orientation using Gabor patches. Subsequently, we examined whether awake reactivation occurs after a brief task on trained and untrained orientations. Our results demonstrate that one of two opponent processes, either awake reactivation or suppression occurs after exposure to untrained and trained orientation, respectively. Immediately after exposure to untrained orientation, the activity patterns of V1 were more likely to be classified as the untrained, compared to trained orientation. In contrast, immediately after exposure to trained orientation, the activity patterns of V1, V2, and V3 were less likely to be classified as the trained, compared to untrained orientation. Further analysis showed that the BOLD signals in V1 decreased significantly after exposure to the trained orientation. The results suggest that suppression occurred after exposure to the trained orientation. In addition, greater learning on the trained orientation was associated with greater reduction of classification as trained orientation. These results demonstrate that the brain operates two opponent processes depending on the novelty of the task during post-learning rest. Exposure to novel task leads to awake reactivation, whereas exposure to extensively trained task leads to awake suppression.

## Results

We investigated if the task’s novelty affects awake reactivation in human. We trained subjects (n = 12) to detect a specific Gabor orientation, either 45° or 135° for two days (**Figure 1**; Please note that 6 subjects who participated in Bang et al. (2018) study volunteered in the current study again. Therefore, they had 1 more day of training on the same orientation from the previous Bang et al. (2018) study compared to other 6 subjects; See Materials and Methods for more details). Subsequently, each orientation was assigned to either trained or untrained orientation. We confirmed that the series of training induced significant learning on the trained orientation (Supplementary figure 1; main effect of time: F(1,11)=7.441, p=0.020, partial η^2^=0.404, two-way repeated ANOVA with factors time (pre vs post) and orientation (trained orientation vs untrained orientation)). The learning amount was not significantly different between trained and untrained orientations, suggesting that learning transferred (interaction between time and orientation: F(1,11)=0.336, p=0.574, partial η^2^=0.030). This generalized learning was observed in our previous and many others’ studies (Bang, Sasaki, et al., 2018; McGovern, Webb, & Peirce, 2012; Wang, Zhang, Klein, Levi, & Yu, 2014; Xiao et al., 2008; Zhang et al., 2010).

**Figure 1.**
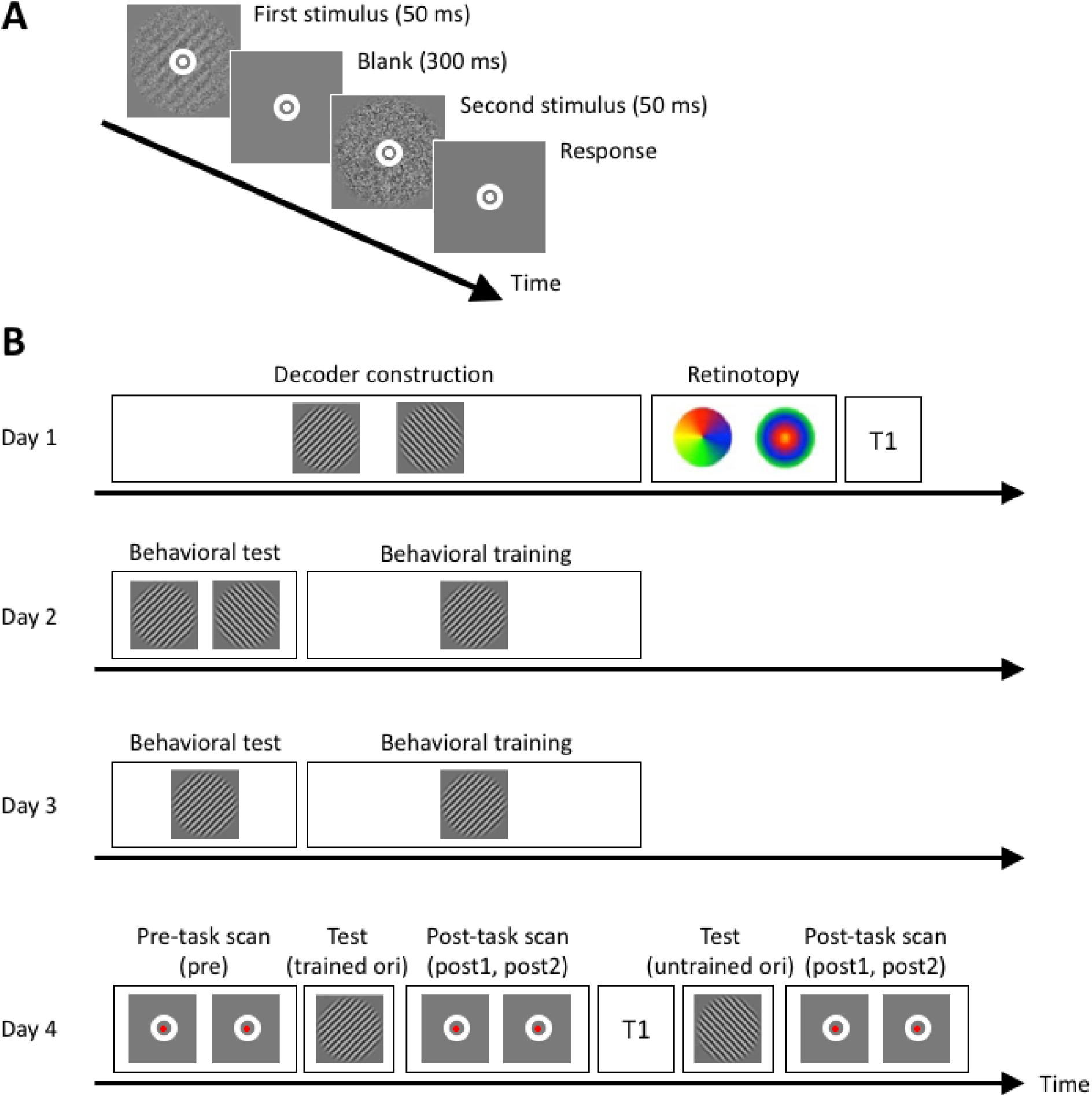
Task and experimental procedure. **A**. Subjects performed 2IFC orientation detection task during behavioral tests and trainings on Days 2 and 3, and tests on Day 4. Two stimuli, a target consisting of a Gabor patch with noise and a non-target consisting of pure noise were presented in a random order. Subjects were required to indicate which interval contained a target. **B**. The experiment consisted of 4 days. Subjects were trained on one orientation among 45° and 135° on Days 2 and 3. On the critical Day 4, We recorded subjects’ brain activity patterns before any task (two 5 min scans; combined into a single ‘pre’), immediately after a task on one orientation (either trained or untrained orientation; post1 and post2), and immediately after a task on the other orientation (either trained or untrained orientation; post1 and post2). The task order of orientations was pseudorandom. Half subjects had trained orientation first, whereas other half subjects had untrained orientation first. During the pre-task and post-task scans, a fixation task was provided to discourage mental imagery of any orientation. In order to examine whether the brain activity patterns within retinotopically-defined visual areas appear similar to a certain orientation, we had a decoder construction scan and a retinotopy scan on Day 1.

We constructed a decoder that could distinguish between trained and untrained orientations for each subject based on the brain activity patterns observed while subjects viewed each Gabor orientation. We applied this decoder to the brain activity scans performed before (pre: two 5 min scans were combined during analysis) and after a brief task on trained and untrained orientations (post1 and post2: consecutive two 5 min scans after each task, the presentation order of two orientations was pseudorandom). Particularly, we focused on early visual areas including V1, V2, and V3 because previous studies showed that plasticity occurs in early visual areas (Rosenthal, Mallik, Caballero-Gaudes, Sereno, & Soto, 2018; Sasaki et al., 2010; Watanabe & Sasaki, 2015).

### Awake reactivation in V1 after exposure to untrained task

First, we examined whether awake reactivation occurs shortly after exposure to the untrained orientation, as similarly shown in our previous study (Bang, Sasaki, et al., 2018). If awake reactivation occurs, the brain activity patterns should appear similar to the untrained compared to the trained orientation after exposure to the untrained orientation. To examine this, we calculated the probability that the decoder would classify the brain activity patterns as untrained orientation. We conducted a two-way repeated measures ANOVA with factors time (pre vs post1 vs post2) and region (V1, V2, V3) to the decoder’s classification. The results revealed a significant interaction between time and region (Figure 2; F(4,44)=3.094, p=0.025, partial η^2^=0.220). Further post-hoc tests showed a significant simple main effect of time in V1 (F(2,10)=17.917, p<0.001, partial η^2^=0.782) but not in V2 (F(2,10)=0.220, p=0.806, partial η^2^=0.042) and V3 (F(2,10)=3.076, p=0.091, partial η^2^=0.381). Additional analyses showed that the probability of untrained orientation became significantly greater immediately after (post1) compared to before (pre) the task in V1 (pre vs post1; T(11)=-3.209, p=0.008, two-sided paired t-test). This increased probability of untrained orientation returned to baseline during the following period, which spanned 5-10 minutes after the offset of the task (pre vs post2; T(11)= - 0.214, p=0.834, two-sided paired t-test). This quadratic trend in V1 was significant, indicating a peak at post1 (F(1,11)=31.082, p<0.001). Furthermore, the classification as untrained orientation was significantly greater than chance shortly after the task (post1: T(11)=3.452, p=0.005, one-sample t-test), but not before (pre: T(11)=-1.581, p=0.142, one-sample t-test) and 5-10 minutes after the task (post2: T(11)=-1.091, p=0.299, one-sample t-test).

**Figure 2.**
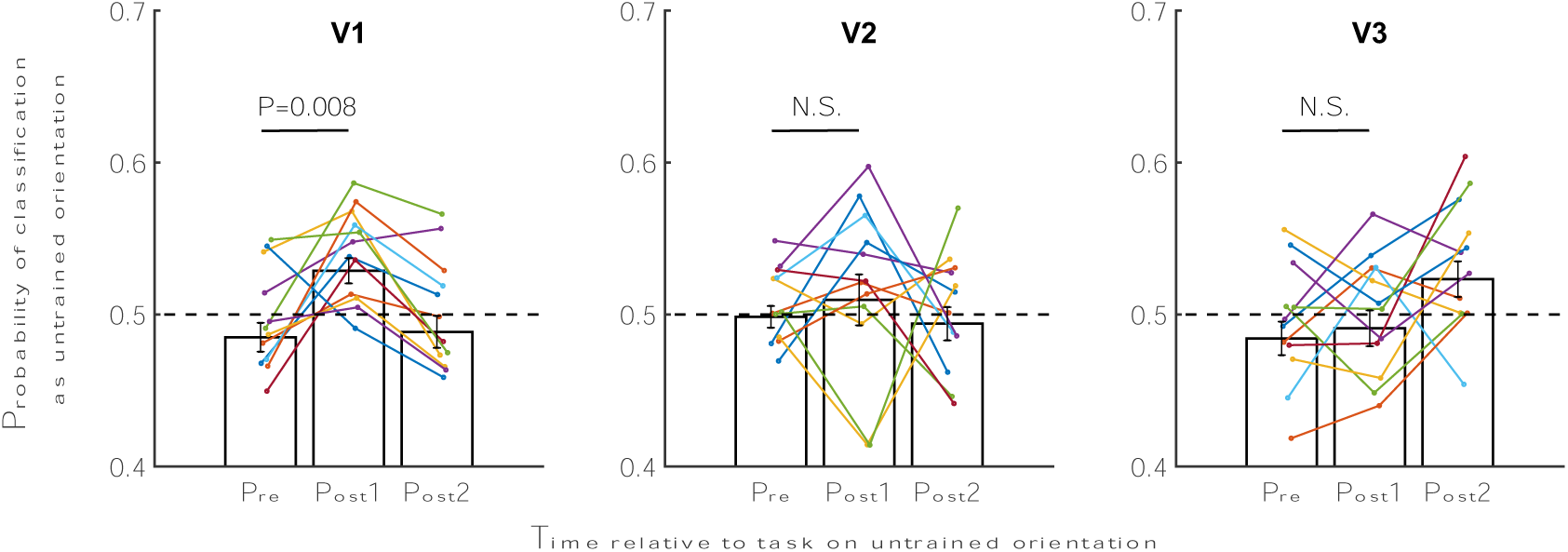
Probability that the brain activity is classified as untrained orientation before and after the task on untrained orientation. The brain activity was more likely classified as untrained orientation in V1 shortly after exposure to untrained orientation. Error bars indicate s.e.m. N.S., not significant.

We also examined whether the task order altered the classification result. Six subjects performed the task on untrained orientation first (order: untrained orientation first, and then trained orientation second) whereas other six subjects performed the task on untrained orientation second (order: trained orientation first, and then untrained orientation second). We found that the probability of untrained orientation shortly after the task was not affected by the order of the tasks in V1 (T(10)=-0.911, p=0.384, independent sample t-test), V2 (T(10)=0.410, p=0.690, independent sample t-test), and V3 (T(10)=-0.958, p=0.361, independent sample t-test). In particular, the increased probability of untrained orientation in V1 was comparable between those who were exposed to untrained orientation first (0.521 ± 0.013 (mean ± s.e.m.)) and second (0.536 ± 0.011 (mean ± s.e.m.)). These results suggest that awake reactivation occurred in V1 shortly after exposure to untrained orientation and was not affected by the task order.

### Suppression in V1, V2, V3 after exposure to trained task

We further examined if exposure to trained orientation, which is of low novelty to the visual system induces awake reactivation as in the case of untrained orientation. For this, we analyzed the decoder’s probability to classify the brain activity patterns as trained orientation. Two-way repeated measures ANOVA with factors time (pre vs post1 vs post2) and region (V1, V2, V3) revealed a significant main effect of time (Figure 3; F(2,22)=5.046, p=0.016, partial η^2^=0.314). Other main effect of region or interaction between time and region were not significant (main effect of region: F(2,22)=1.801, p=0.189, partial η^2^=0.141; interaction between time and region: F(4,44)=0.329, Huynh-Feldt correction, p=0.789, partial η^2^=0.029). Additional post-hoc tests showed that the probability of trained orientation became significantly reduced after (post1) compared to before (pre) the task (pre vs post1; p=0.013, 95% CI=0.002−0.067). Reduced probability returned to baseline 5-10 minutes after the offset of the task (pre vs post2: p=0.333, 95% CI=-0.020−0.040). This reduction of the probability was not altered by the order of task. The classification as trained orientation shortly after the task (post1) was not significantly different between those who were exposed to the trained orientation first and second (V1: T(10)=-0.363, p=0.724, V2: T(10)=0.106, p=0.918, V3: T(10)=0.694, p=0.504, independent sample t-test). The probability of trained orientation immediately after the task dropped to 0.474 ± 0.018 (mean ± s.e.m.) and 0.476 ± 0.034 (mean ± s.e.m.) for those who were exposed to the trained orientation first and second, respectively. Apparently, this dip in the probability after exposure to trained orientation is in contrast to enhanced probability after exposure to untrained orientation.

**Figure 3.**
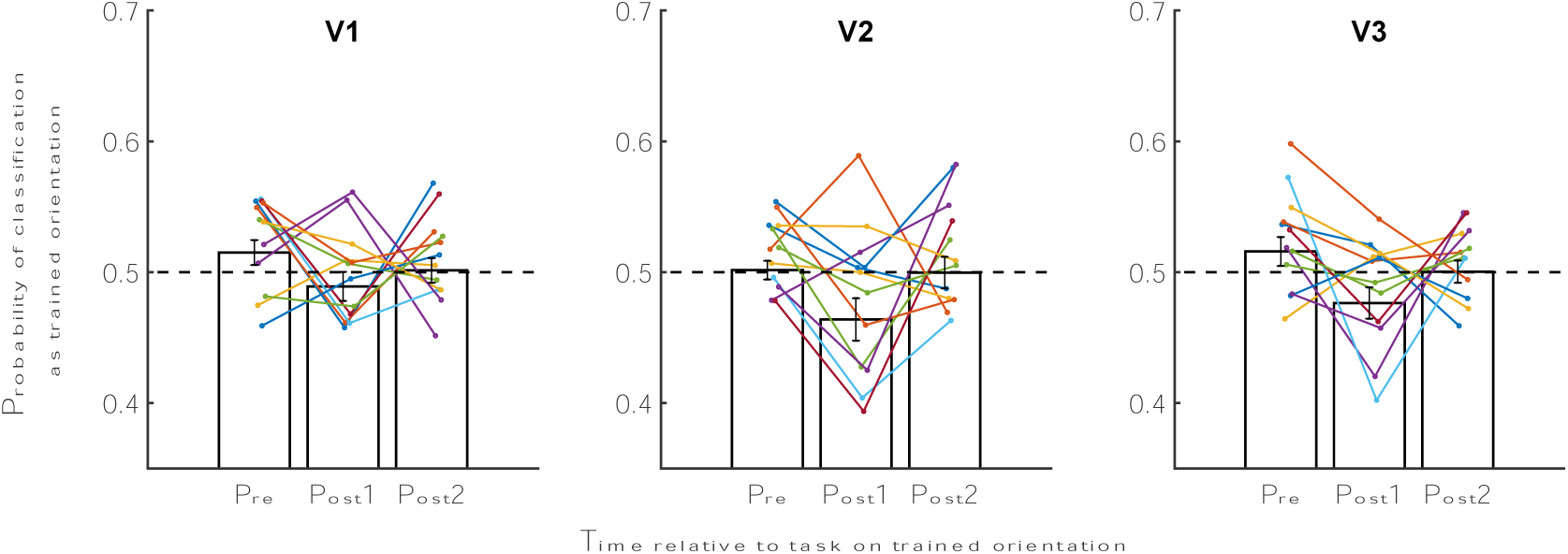
Probability that the brain activity is classified as trained orientation before and after the task on trained orientation. The brain activity appeared more similar to trained orientation in V1, V2, and V3 shortly after exposure to trained orientation. Error bars indicate s.e.m. * p<0.05. N.S., not significant.

We then examined if decreased classification as trained orientation reflects suppression of trained orientation in early visual areas. We reasoned that if suppression occurred, the voxels that encode trained orientation would demonstrate decreased BOLD signals after (post1) compared to before the task (pre). To examine this, we calculated the amplitude index for trained orientation for each subject (for details about the index, see Methods). We then examined if the amplitude index for trained orientation changed immediately after the task in V1, V2, and V3. The results showed a significant decrease of the amplitude index in V1 (Figure 4; T(11)=2.732, p=0.020, paired sample t-test). We did not find such changes in V2 (T(11)= - 1.464, p=0.171) and V3 (T(11)=0.470, p=0.648), although the decreased classification as trained orientation was observed in all V1, V2, and V3. The amplitude results suggest that the representation of trained orientation was suppressed in V1 after exposure to the trained orientation.

**Figure 4.**
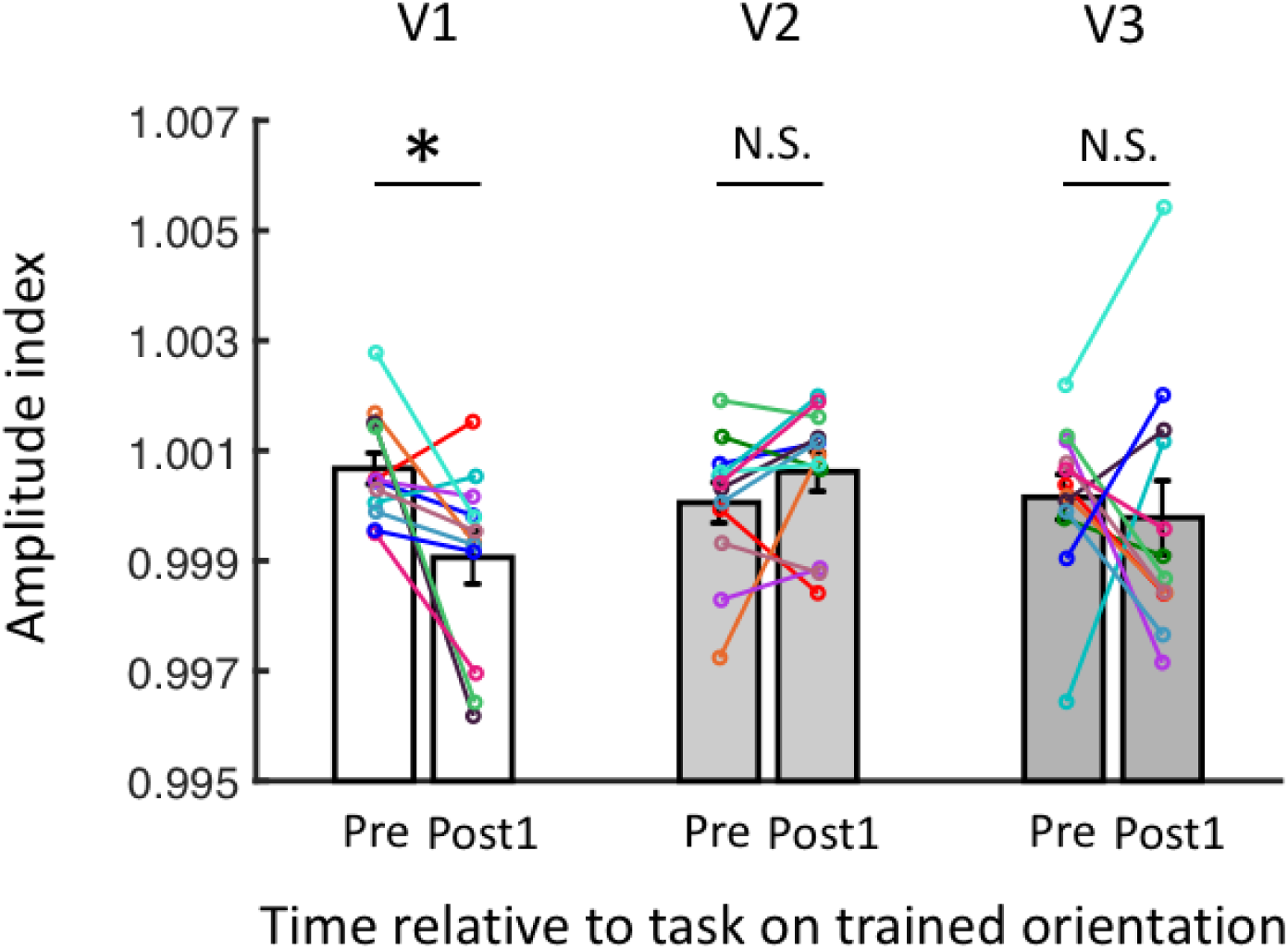
Reduced amplitude index in V1 immediately after exposure to trained orientation. The amplitude index (mean ± s.e.m.) in V1 significantly reduced shortly after a task on trained orientation. Such reduction was not found in V2 and V3. * p<0.05. N.S., not significant.

### Impact of task novelty on post-task classification

Having analyzed the post-task processes separately for the untrained and trained orientations, we further tested the effect of the task novelty by entering both data into a three-way repeated measures ANOVA. For this, we calculated the probability that the decoder would classify the brain activity patterns as the exposed orientation. Then, we performed a three-way repeated measures ANOVA with factors task type (untrained vs trained orientation), time (pre vs post1 vs post2) and region (V1, V2, V3) on the decoder’s probability of the exposed orientation. If there is an impact of task novelty on the post-task processes, the probability that the brain patterns appear similar to the exposed orientation should be different depending on whether the task involved trained or untrained orientation. The results showed a significant interaction between task type and time (F(2,22)=4.705, p=0.020, partial η^2^=0.300). Further post-hoc tests showed a significant simple main effect of task type immediately after the task (post1: first 5 min scan after the task; F(1,11)=5.994, p=0.032, partial η^2^=0.353) but not before (pre: two 5 min scans before the task; F(1,11)=2.392, p=0.150, partial η^2^=0.179) and 5 minutes after the task (post2: second 5 min scan after the task; F(1,11)=0.038, p=0.848, partial η^2^=0.003). Immediately after the task (post1), the probability of exposed orientation was significantly greater for the task involving untrained orientation compared to the task involving trained orientation (p=0.032, 95% CI=0.003−0.063). These results are consistent with that awake reactivation occurs after exposure to the untrained orientation, but awake suppression occurs after exposure to the trained orientation.

### No awake reactivation or suppression outside the early visual areas

We expanded our analysis to the whole brain to investigate if any changes in the classification occurred after the task. We created anatomical ROIs encompassing subareas of the frontal, temporal, and parietal areas. We also included retinotopically defined higher visual areas such as V3A, and V4v (see Methods). We performed leave-one-run-out cross-validation (10-fold cross-validation) to test which ROIs contain orientation-specific information. The results showed significant accuracy in five ROIs: fusiform area (T(11)=2.231, p=0.047), middle temporal cortex (T(11)=2.794, p=0.017), superior parietal cortex (T(11)=3.790, p=0.003), V3A (T(11)=7.154, p<0.001), and V4v (T(11)=6.047, p<0.001). Based on the cross-validation results, we further examined if the classification as the exposed orientation changed after task on either untrained and trained orientation in each of these ROIs. A one-way repeated ANOVA with a factor time (pre vs post1 vs post2) showed no significant main effect of time in any of ROIs (all p values > 0.1). Although we did not find any overall effect of time, we further tested if the probability of the exposed orientation changed immediately after compared to before the task. No ROIs showed such changes in the probability of the exposed orientation (all p values > 0.1). These results suggest that previously observed awake reactivation and suppression were confined in early visual areas.

### Greater learning is associated with greater suppression

Finally, we examined the relationship between behavioral performance and classification changes. Specifically, we examined if performance improvement on the trained orientation is associated with reduced probability of trained orientation after exposure to trained orientation. For this, we constructed individual’s learning index for trained orientation (for more details about learning index, see Methods). Then we compared if the reduced probability of the trained orientation immediately after the task was significantly different between those who had higher versus low amount of learning using a median split. The results showed that those who had higher performance improvement had greater reduction in the probability of trained orientation than those who had lower performance improvement (Figure 5; T(10)=-2.425, p=0.036). This association suggests that the suppression amount of the trained feature depends on how much performance improvement the individual obtained. In short, the greater the learning is, the greater the suppression is.

**Figure 5.**
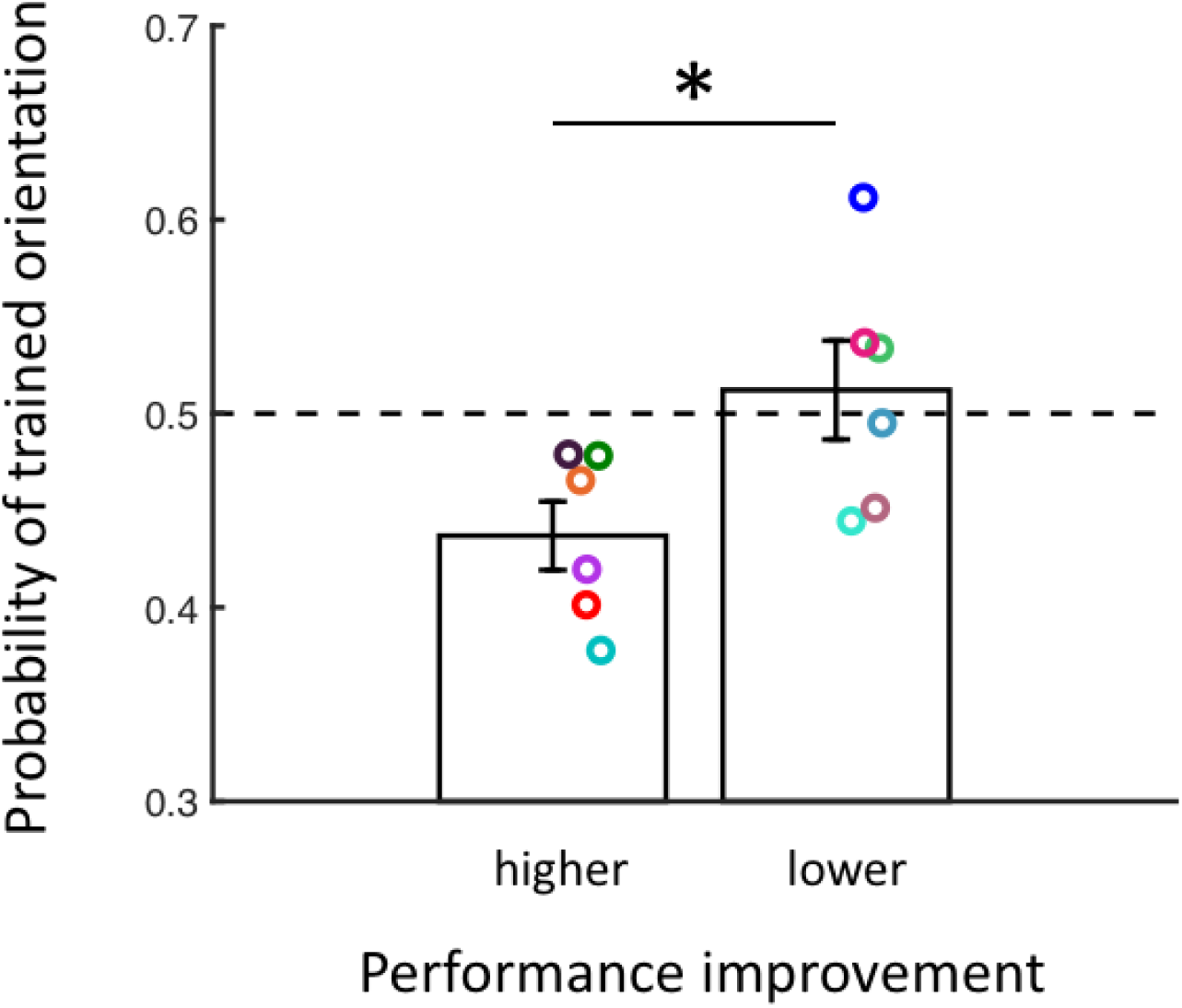
Greater learning is associated with greater suppression. The probability of trained orientation (mean ± s.e.m.) was significantly lower for those who showed greater learning on the trained orientation based on a median split. Dots represent individual data. * p<0.05.

## Discussion

We examined how the novelty of the task affects post-task neural processes in human early visual areas. We found that the brain operates two opponent processes, either awake reactivation or suppression after a brief task on untrained and trained orientations, respectively. The awake reactivation was observed in V1, whereas the suppression appeared in early visual areas including V1, V2, and V3. Additionally, we observed that the performance improvement is associated with the suppression amount. Those who improved more showed greater suppression of the trained feature. These results provide clear evidence that different post-task processes occur depending on the task novelty.

Accumulating studies have shown that newly encoded information is reactivated during the following rest period (Bang, Sasaki, et al., 2018; Chelaru et al., 2016; de Voogd et al., 2016; Deuker et al., 2013; Guidotti et al., 2015; Schlichting & Preston, 2014; Staresina et al., 2013; Tambini & Davachi, 2013). This awake reactivation has been observed to play a key role in consolidating memories. Although many studies have examined the nature of awake reactivation, relatively little is known about how this awake reactivation is modulated by other task-related factors. The current study aimed to examine the effect of the task novelty on awake reaction. The results show that awake reactivation occurs when the orientation is novel but suppression, the opposite of awake reactivation, occurs when the orientation has little novelty. We posit that these opponent processes, awake reactivation or suppression after exposure to task of high or low novelty are in line with previous findings. Previous studies showed that information paired with reward (Gruber, Ritchey, Wang, Doss, & Ranganath, 2016) or shock (de Voogd et al., 2016) was preferentially reactivated during following rest. It is highly likely that reward and shock made the paired information more salient than other information and then the brain selected salient information to be reactivated more in the following rest. Even without such external modulator, it seems that the brain has its own selection mechanism. Recent study showed that the brain prioritizes weakly learned information and predicts later performance (Schapiro, McDevitt, Rogers, Mednick, & Norman, 2018). Together with this line of studies, the current study adds the evidence that the reactivation favors novel and salient information.

Critically, suppression of exposed feature shortly after task suggests that inhibition process is involved in the post-task processes. Pattern suppression was previously observed as a forgetting mechanism of competing memories while target memories were retrieved (Wimber, Alink, Charest, Kriegeskorte, & Anderson, 2015). However, in the post-task processes, such suppression has not been observed to our best knowledge. Therefore, there remain much questions about how the suppression affects memory, and what functions the suppression performs. Although not directly related, recent study showed that overlearning on the visual task (same 2IFC orientation detection task was used) changes the neurochemical environment of the early visual areas into inhibitory-dominant state shortly after task (Shibata et al., 2017). Such inhibitory-dominant state was found to be associated with hyperstabilization of visual learning. Hyperstabilization is the state where the learning becomes not only resilient against interference from the following new learning, but also disrupts the new learning. Additionally, GABA was shown to be associated with BOLD signals negatively (Northoff et al., 2007). Therefore, there could be some common mechanisms between the current study’s suppression after trained orientation and inhibitory-dominant state after overlearning.

Similarly, it remains to be seen whether the current study’s awake reactivation after untrained orientation is related with excitatory-dominant state after visual task. Previous studies showed that the early visual area’s neurochemical environment changed to an excitatory-dominant state after offset of visual task (same 2IFC orientation detection task was used) (Bang, Shibata, et al., 2018; Shibata et al., 2017). In particular, one study used 8 blocks of visual task (Shibata et al., 2017) and the other study used 3 blocks of visual task that was trained on the previous day (Bang, Shibata, et al., 2018). In our study, we used 3 blocks of visual task. Given that these studies used the same task with similar number of blocks, there could be some common mechanisms between the current study’s awake reactivation and excitatory-dominant state after visual task. One may think 3 blocks of visual task that was trained on the previous day is similar to our 3 blocks of visual task on trained orientation. However, it needs to be noted that the previous study had 1 day of training, whereas our study had 2-3 days of training. Since literatures have reported that visual plasticity involves different dynamics between early and late phase of learning, much caution is needed when inferring plasticity from different days of training (Kang et al., 2018; Yotsumoto, Watanabe, & Sasaki, 2008). It also remains unexplored yet how these awake reactivation and excitatory-dominant state after visual task are linked to the plastic state of visual learning. Two above-mentioned studies showed that excitatory-dominant state was associated with the plastic state of visual learning. In other words, visual learning was vulnerable against interference during excitatory-dominant state. Therefore, our study casts a question whether awake reactivation is associated with instability of visual learning.

In conclusion, we found that opposite post-task processes occur depending on the task novelty. Awake reactivation occurred after exposure to an untrained orientation, whereas awake suppression occurred after exposure to a trained orientation. Furthermore, previous learning amount was positively associated with the strength of suppression. These results suggest that awake suppression and awake reactivation might be critical offline mechanisms that are functionally related to visual learning.

## Materials and Methods

### Participants

Twelve subjects (19-27 years old, 5 females) participated in this study. All subjects had normal or corrected-to-normal vision and did not have any history of neurological disorders. All subjects were screened for MRI safety and provided written informed consent. The study was approved by Institutional Review Board of Georgia Institute of Technology. Six among twelve subjects previously participated in Bang et al. (2018)’s study. The sample size was determined based on previous fMRI experiments on visual perceptual learning (Bang, Sasaki, et al., 2018; Guidotti et al., 2015; Shibata, Sasaki, Kawato, & Watanabe, 2016).

### Task

Subjects performed 2-interval-forced-choice (2IFC) orientation detection task. Subjects were provided with a target and non-target stimuli in a random order and indicated which of two intervals contained a target. The target was a Gabor patch containing a particular orientation (circular diameter = 10 degree, contrast = 100%, spatial frequency = 1 cycle/degree, Gaussian filter = 2.5 degree, random spatial phase). The Gabor patch was masked by noise generated from a sinusoidal luminance. The stimulus intensity was controlled by the ratio of noise pixels. For example, X% S/N ratio contained noise in 100-X% of the pixels in the Gabor patch. The other non-target stimulus consisted of noise only (0% S/N ratio).

Each trial consisted of a 500-ms fixation period and two intervals containing the target and non-target stimuli in a random order lasting for 50 ms each. Two intervals were separated by a 300-ms blank period. After two intervals, subjects reported which interval had the target by pressing a button on a keypad under no time restriction. No feedback was given after the response.

### Procedures

The study consisted of 4 separate days (see Figure 1). On Day 1, subjects had decoder construction, retinotopy and anatomical MRI scans. On Day 2, subjects had pre-training behavioral tests on two orientations and training on one randomly chosen orientation among two. Day 3 consisted of behavioral test and training on the trained orientation. On Day 4, subjects had pre-task scan, behavioral test on one orientation (either trained or untrained orientation) and immediately following post-task scan. Then subjects had T1 anatomical scan followed by behavioral test on the other orientation (either trained or untrained orientation) and immediately following post-task scan. Days 1, 2, and 3 could be separated by multiple days, but Days 3 and 4 were constrained to be separated up to 3 days. Days 1 and 4 were conducted in 3T scanner, but Days 2 and 3 were performed in a mock scanner.

The behavioral test and training consisted of 3 and 10 blocks, respectively. Each block began with 25% S/N ratio and the following S/N ratios were determined by a 2-down-1-up staircase procedure. The block terminated after 10 reversals. On average, each block had 30-40 trials and took 1-2 minutes.

The behavioral test on Day 2 was performed for each 45° and 135° orientation. The order of two orientations was determined randomly. Once determined, subjects performed 3 blocks for the first orientation, and then another 3 blocks for the second orientation. The purpose of the behavioral test was to measure individual’s threshold S/N ratio for each orientation. The threshold S/N ratio per block was calculated as the geometric mean of the last 6 reversals, following previous studies (Bang, Milton, Sasaki, Watanabe, & Rahnev, 2019; Bang, Sasaki, et al., 2018; Bang, Shibata, et al., 2018; Shibata et al., 2017). The behavioral training was performed for one randomly chosen orientation among 45° and 135°. The purpose of the behavioral training was to train subjects on one orientation, thus differentiating the novelty of the task between trained and untrained orientations.

Please note that Day 1 experiments, including decoder construction, retinotopy and anatomical MRI scans overlap with previous Bang et al. (2018)’s Day 1 experiments. 6 subjects who previously participated in Bang et al. (2018) volunteered in the current study. Therefore, these 6 subjects participated from Day 2 experiments in the current study. Their existing Day 1 data from Bang et al. (2018) were reused in the current study. In addition, please note that these 6 subjects were previously trained on either 45° or 135° orientation from Bang et al. (2018)’s Day 2 experiments. To maintain consistence in training, we trained these 6 subjects on the same orientation where they were previously trained.

### Decoder construction

In order to construct the decoder that can distinguish between two Gabor orientations, we collected individual’s BOLD signal patterns while subjects viewed each Gabor orientation (45° and 135°). For this, subjects performed frequency detection task (the same task was used in Bang et al. (2018)). Subjects were presented with series of Gabor patches with orientations (either 45° or 135°) and determined whether there was a change in the spatial frequency among presented Gabor patches while fixing their eyes at the central dot. The task had 10 runs (1 run = 300 s) each consisting of 18 trials (1 trial = 16 s) with two fixation periods (each 6 s) each in the beginning and the end of the run. Each trial contained 12-s stimulus presentation period and the following 4-s response period. During the stimulus presentation period, 12 Gabor patches with one identical orientation (45° or 135°) flashed at a rate of 1 Hz. Each Gabor patch was presented for 500 ms and there was 500-ms blank period between Gabor patches. All 12 Gabor patches had the same orientation.

In half of trials, 1 among 12 Gabor patches had a higher spatial frequency than other 11 Gabor patches. Subjects were asked to press a button during 4-s response period if they detected a change in the spatial frequency among 12 Gabor patches. In the beginning of the 12-s stimulus presentation period, the central dot changed its color from white to green. This dot remained green during the entire 12-s stimulus presentation period, and then returned to white when 4-s response period began.

The difficulty of the task was controlled by the degree of the spatial frequency change. The first spatial frequency change in the first run was fixed to 0.24 cycles/degree and the following changes were adjusted via adaptive staircase method. For the case of hit or miss, the spatial frequency change decreased or increased by 0.02, respectively. When a correct rejection was made, the frequency did not change. The following run’s spatial frequency started from the previous run’s last spatial frequency.

The order of orientations across 18 trials per run was pseudorandom in a way that half trials contained 45° orientation and the other half had 135° orientation.

### Pre-task and post-task scans

The purpose of the pre-task and post-task scans was to obtain brain activity patterns during before and after the task on untrained and trained orientations. The pre-task scan was performed first before any task was given, and then the post-task scan was performed twice, one after the task on one orientation and then another after the task on the other orientation. The pre-task and post-task scans involved two 5-min scans for each. In the analysis, we combined two 5-min scans of the pre-task scan into one because we did not expect a change during two 5-min scans. On the other hand, we analyzed two 5-min scans of the post-task scan separately (post1, post2) to examine the time course changes of post-task processes.

During the pre-task and post-task scans, subjects performed a fixation task (the same task was used in Bang et al. (2018)). It was designed to discourage subjects from imagining a certain orientation during scans. Subjects were asked to maintain their fixation at the central dot (size and location of the dot was identical with one used in the 2IFC task) and press a button when they detected a color change of the dot. The central dot’s color changed from white to faint pink ([R, G, B] = [255, 255 – x, 255 – x]) for 1.5 s and then returned to white. Subjects’ response had to be made within this 1.5 s. The difficulty of the task was controlled by the color change. The color change x was set to 40 in the beginning and then adjusted via 2-down 1-up staircase procedure. The step size was fixed to 2.

### MRI data acquisition

Subjects’ MRI data were collected in a Siemens 3T Trio MR scanner using a 12-channle head coil. Anatomical images were obtained using a T1-weighted MPRAGE sequence (256 slices, voxel size = 1 × 1 × 1 mm^3^, TR = 2530 ms, FOV = 256 mm). Functional images were collected using a gradient echo-planar imaging sequence (33 slices, voxel size = 3 × 3 × 3.5 mm^3^, TR = 2000 ms, TE = 30 ms, flip angle = 79°). The slices covered the whole brain and were parallel to the AC-PC plane.

### Retinotopy and ROI selection

Using standard retinotopy method (Sereno et al., 1995; Tootell et al., 1997), we presented a flickering checkerboard along the horizontal and vertical meridians and in the upper and lower visual fields. Using contrast maps between horizontal versus vertical meridians and upper versus lower visual fields, we delineated V1, V2, V3, V3A, and ventral V4. We also obtained 12 anatomical ROIs using Freesurfer’s cortical parcellation method. These ROIs include orbitofrontal cortex, superior, middle and inferior frontal cortex, precentral, postcentral, and paracentral cortex, superior and inferior parietal cortex, and superior, middle, and inferior temporal cortex.

### fMRI data analysis

We used Freesurfer software to analyze the MRI data. First, we used the longitudinal stream in Feesurfer (Reuter, Schmansky, Rosas, & Fischl, 2012) to extract reliable structural images from two different days’ scans (Days 1 and 4). This method generates an unbiased within-subject template using robust, inverse consistent registration. The functional data from two different days were preprocessed using Freesurfer. We applied motion correction, but not spatial or temporal smoothing on the functional data. The functional images were registered to the individual structural template that was created by the longitudinal stream. Then we used a gray matter mask to extract BOLD signals from the voxels located within the gray matter.

Using Matlab, we shifted the BOLD signals by 6 s to account for the hemodynamic delay. We removed voxels that had spikes greater than 10 SDs from the mean during decoder construction scan. We also removed a linear trend in the time course BOLD signals. The BOLD signals in each run were then normalized to z-score for each voxel. We averaged the BOLD signals of each voxel by every 6 volumes (12 s) to create the data sample for decoding. We obtained 90 data samples for each orientation, thus in total 180 data samples from 10 runs of decoder construction scan. Of note, 6 volumes correspond to the Gabor stimulus presentation time during the decoder construction scan.

We used linear sparse logistic regression for decoding. This method selects the relevant voxels in the ROIs and calculates their weight parameters for classification. We trained the decoder to classify the brain activity patterns in each ROI to either 45° or 135° using 180 data samples. The decoder’s reliability was tested using a 10-fold cross-validation. The decoder was trained on nine runs and tested on the remaining run. We observed a significant performance for all retinotopically defined ROIs (V1: T(11)=8.587, p<0.001; V2: T(11)=10.327, p<0.001; V3: T(11)=7.287, p<0.001; V3A: T(11)=7.154, p<0.001; V4v: T(11)=6.047, p<0.001; uncorrected one-sample t-tests).

Based on the high performance of the decoder, we further applied the decoder to the pre-task and post-task scans. We used the same voxels that were included during the decoder construction. We shifted the BOLD signals by 6 s, removed a linear trend, z-scored and then averaged by every 6 volumes. Then we applied the decoder to each 6-volume period of the pre-task and post-task scans. The decoder calculated how probable each 6-volume data sample represents the untrained or trained orientation. We averaged these probability scores within each pre, post1, and post2 to obtain the decoder’s probability of classifying brain activity as the untrained and trained orientation.

### Statistics

For all statistical tests, we used two-tailed parametric tests. We used Mauchly’s Sphericity test for all repeated measures ANOVAs to test the assumption of sphericity. When the sphericity assumption was violated, we used Huynh-Feldt correction. We reported all such violations.

### Amplitude index

In order to examine if the BOLD signals are suppressed during post-task scan shortly after exposure to trained orientation, we calculated each individual’s amplitude index. The amplitude index was defined as the mean amplitude of the diagnostic voxels for the trained orientation compared to the mean amplitude of the remaining non-diagnostic voxels. For this, we first selected diagnostic voxels for the trained orientations that have positive weight for the trained orientation. We calculated the mean amplitude of these diagnostic voxels for the trained orientation. Then we divided it by the mean amplitude of non-diagnostic voxels that have zero weight for the trained orientation. This amplitude index allows us to track the relative amplitude of the diagnostic voxels compared to the remaining other non-diagnostic voxels.

### Learning index

We defined each individual’s learning index as the relative performance improvement between before (Day 2’s test) and after training (Day 4’s test). We measured the threshold S/N ratio per session as the mean threshold S/N ratios across three blocks. We subtracted the threshold S/N ratio after training (Day 4’s test) from that before training (Day 2’s test) and then divided it by the threshold S/N ratio before training (Day 2’s test) to obtain the learning index.

### Apparatus

We created all visual stimuli in Matlab using Psychophysics Toolbox 3. We used LCD display (1024 × 768 resolution, 60Hz refresh rate) inside a mock scanner and MRI-compatible LCD projector (1024 × 768 resolution, 60Hz refresh rate) inside the 3T scanner to present the visual stimuli.

---

**Supplementary Figure 1.**
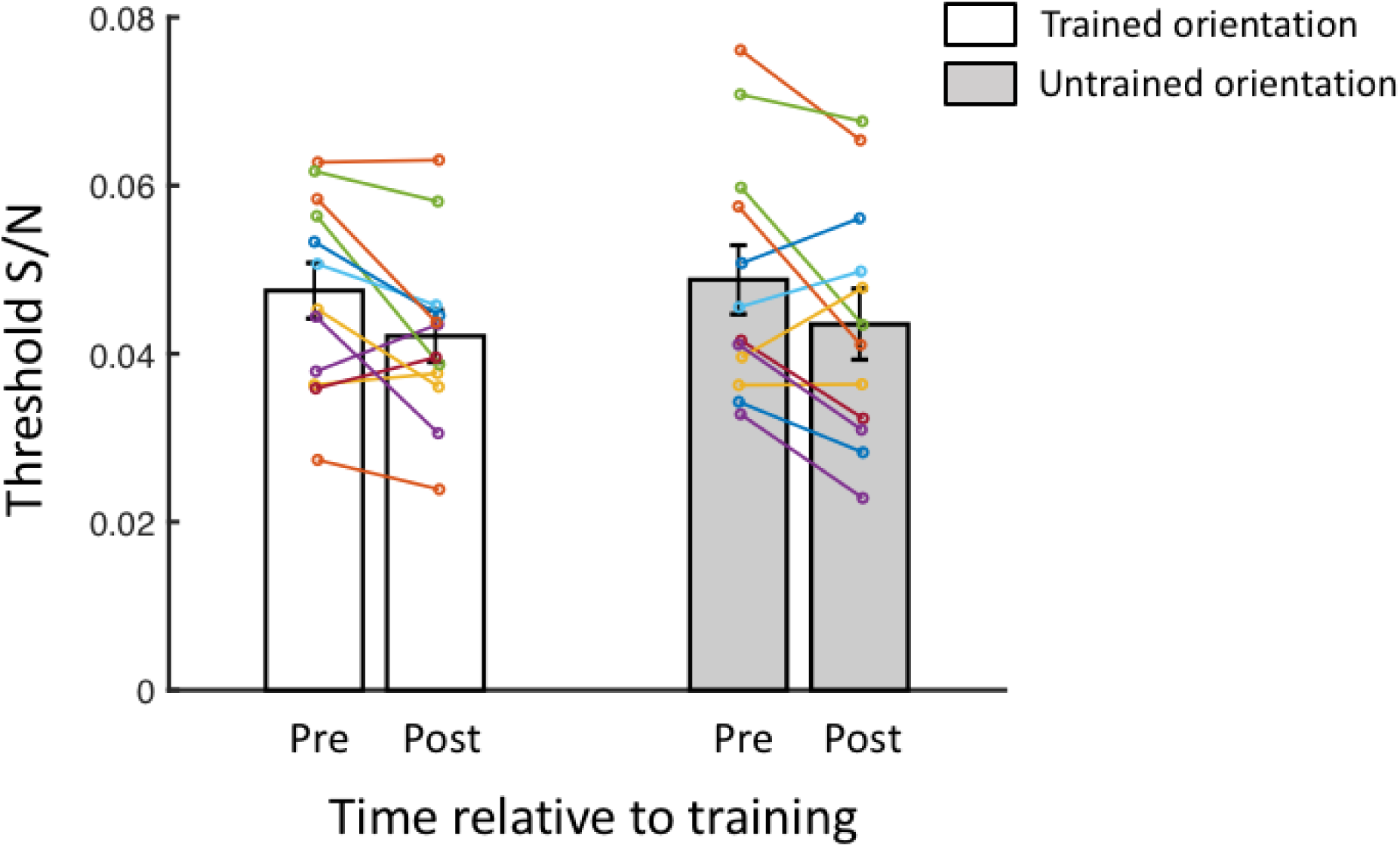
Threshold S/N (mean ± s.e.m.) for the trained and untrained orientations before (Day 2’s behavioral test) and after training (Day 4’s Test). To examine if learning occurred after training, we performed a two-way repeated ANOVA with factors time (pre vs post) and orientation (trained orientation vs untrained orientation). The results showed a significant main effect of time (F(1,11)=7.441, p=0.020, partial η^2^=0.404), suggesting that learning occurred after training. However, the learning amount did not differ between trained and untrained orientations (interaction between time and orientation: F(1,11)=0.336, p=0.574, partial η^2^=0.030). It is likely that this learning transfer was driven partly by exposure to both orientations during decoder construction scan on Day 1. The dots indicate individual data.

**Supplementary Figure 2.**
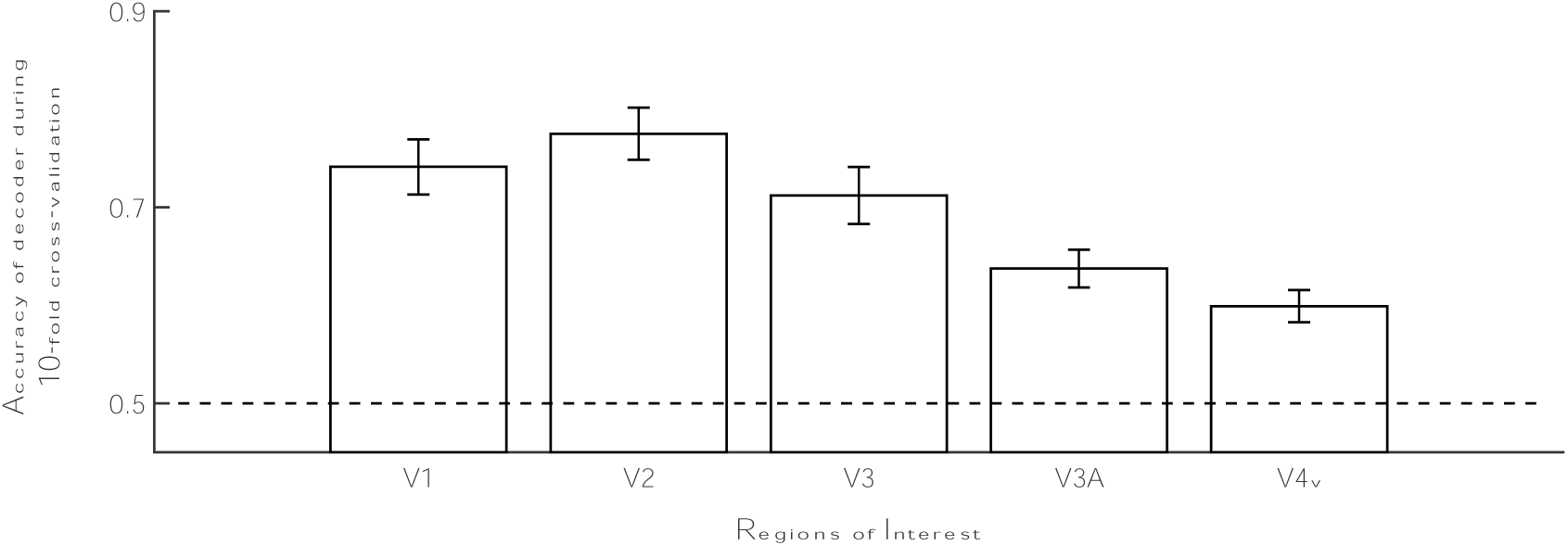
Accuracy of decoder during 10-fold cross-validations in V1, V2, V3, V3A, and V4v. The accuracy of decoder (mean ± s.e.m.) was significantly greater than chance for all these areas (V1: T(11)=8.587, p<0.001; V2: T(11)=10.327, p<0.001; V3: T(11)=7.287, p<0.001; V3A: T(11)=7.154, p<0.001; V4v: T(11)=6.047, p<0.001; uncorrected one-sample t-tests).

